# Natural variation at the *Drosophila melanogaster Or22* odorant receptor locus is associated with changes in olfactory behaviour

**DOI:** 10.1101/2021.05.27.446061

**Authors:** Katherine H. Shaw, Craig I. Dent, Travis K. Johnson, Alisha Anderson, Marien de Bruyne, Coral G. Warr

## Abstract

In insects many critical olfactory behaviours are mediated by the large odorant receptor (Or) gene family, which determine the response properties of different classes of olfactory receptor neurons (ORNs). While ORN responses are generally conserved within and between Drosophila species, variant alleles of the *D.melanogaster Or22* locus have previously been shown to the response profiles of an ORN class called ab3A. These alleles show potential clinal variation, suggesting that selection is acting at this locus. Here, we investigated if the changes seen in ab3A responses lead to changes in olfactory-related behaviours. We show that variation at the *Or22* locus and in the ab3A neurons are not fully compensated for by other ORNs and lead to overall changes in antennal odorant detection. We further show that this correlates with differences in odorant preference behaviour and with differences in oviposition site preference, with flies that have the chimaeric short allele strongly preferring to oviposit on banana. These findings indicate that variation at the *Or22* locus leads to changes in olfactory-driven behaviours that could be under selective pressure, and add support to the idea that the ab3A neurons are of especial importance to the ecology of Drosophila flies.

## Introduction

Animals rely on their sense of smell to discriminate between odours in order to identify and locate mates, dangers and food sources. Flying insects need to be able to do all of this rapidly, and additionally utilise their chemosensory systems to identify the optimal places to lay their eggs. In insects odour identity is combinatorially encoded through the inputs from a large number of classes of olfactory receptor neurons (ORNs) (1–3), whose responses are determined by the olfactory receptors they express (4–6). One of the major families of olfactory receptors in insects is the rapidly evolving Or gene family, which encode highly divergent ligand-gated ion channels and primarily detect volatile odorants (7). The odour responses of this family and their mapping to individual ORN classes have been best characterised in *Drosophila melanogaster*, where there are 62 Ors. In *D. melanogaster* individual Ors/ORNs can be either broadly tuned (responding strongly to a wide range of odorants) or narrowly tuned (responding strongly to one or a few odorants), albeit with there being a continuum in the breadth of tuning rather than a strict dichotomy (8, 9).

We previously reported one of the first described cases of within-species variation in responses of a *Drosophila melanogaster* ORN, the ab3A neuron (10). Among many ORN classes, this ORN is also one of only a few known to show interspecies variation between closely-related Drosophila species (11, 12). The olfactory receptor locus that determines ab3A response is the *Or22* locus, and we showed that naturally occurring variation at this locus within *D. melanogaster* causes alterations to ab3A neuronal responses. Specifically, the *Or22* locus variant found in the common laboratory strain Canton S has two genes encoding Ors at this locus, *Or22a* and *Or22b* (the long allele; [10, 13]), however the Or22b protein is not functional. In these flies Or22a produces an ab3A response phenotype we call ab3A-1. A major variant at this locus that has been found in wild populations (14, 15) is a short allele in which a chimaeric receptor called Or22ab produces a different odour response profile that we call ab3A-2. We also identified a third phenotype with yet another different odour response profile (ab3A-3), in which there is still a long allele (with two gene copies) but *Or22a* is not expressed and instead a functional version of *Or22b* determines ab3A response (10). In general, ab3A-1 neurons respond most strongly to the larger esters tested (e.g. ethyl hexanoate, methyl hexanoate and ethyl octanoate), ab3A-2 neurons to small acetate esters (e.g. propyl acetate, butyl acetate and pentyl acetate) and ab3A-3 neurons to smaller ethyl esters (e.g. ethyl propionate, ethyl butanoate and ethyl 2-methylbutanoate).

Frequencies of the long and short alleles at this locus appear to be clinally varying in Australia, with the long allele being fully penetrant in the south and the short allele appearing at high frequency in the north (15). This clinal variation strongly suggests that selection is acting at this locus, and we therefore wondered if the *Or22* variants cause changes in behaviours that are important for individual fitness. However, because many other neuron types also detect the odorants to which ab3A responds, it remained unknown whether changes at the level of the ab3A neuron affect behavioural responses to odorants. Here, we have investigated this question. We first show that broader antennal neuronal responses to odours are impacted by the ab3A response variation. We further provide evidence of an association between ab3A phenotype and behavioural responses in two different olfactory behaviour paradigms. Taken together, our data strongly suggests that variation at the *Or22* locus leads to changes in overall odorant detection by the antenna that are associated with changes in biologically-significant behaviours upon which selection might act.

## Results

### Variation in ab3A phenotype alters overall antennal detection of ab3A ligands

The ab3A neuron is broadly-tuned, with significant responses (greater than 50 spikes/sec) described to at least 50 odorants (8). Most, if not all, of these are also detected by other ORNs, for example pentyl acetate is also detected by ab1A, ab5B, ab6A and ab7A neurons, and ethyl butanoate is also detected by ab1A, ab8A, ab8B, ab10A, ab3B and ab2B neurons (1, 8). It is therefore possible that changes in detection of odorants by ab3A neurons, for example loss of sensitivity, could be compensated for by changes in other neurons, and in this scenario the ab3A change would seem less likely to impact behaviour. In order to determine if there are differences in broader antennal responses to odorants between the ab3A phenotypes we therefore measured responses from populations of neurons across the antenna using electroantennogram recordings (EAGs). Rather than measuring the response of a single neuron, an EAG signal represents the summed responses of all ORNs in the vicinity of the recording electrode (16) (see (1, 17) for diagram of ORN distribution on the antenna). If compensatory changes are occurring in other ORN classes then we would expect no difference in EAG responses to ab3A-detected odorants in flies with the different ab3A phenotypes. Conversely a change in EAG response would indicate either a partial or complete lack of compensation.

We therefore tested the EAG responses of fly lines isogenic (fully homozygous) for the second chromosome that individually exhibit the ab3A-1, ab3A-2, and ab3A-3 phenotypes to serial dilutions of odorants that we previously showed were able to distinguish the three phenotypes at the single neuron level (10). Odorant concentrations were selected based on our previous data on the ab3A responses to serial dilutions of these odorants, and we selected the lowest dose that gave the maximum response from the relevant phenotype and one dose either side of this. For ethyl hexanoate and ethyl butanoate this gave a range from 10^-4^ to 10^-2^, while for isopentyl acetate this gave a range from 10^-5^ to 10^-3^. Paraffin oil was also tested as the solvent control.

We found significant differences in EAG responses between flies with the different ab3A phenotypes (Figure 1; Supplementary Table 1). Flies with the ab3A-1 phenotype show significantly higher EAG responses to the ab3A-1 ligand ethyl hexanoate at 10^-4^ and 10^-3^ than do ab3A-2 flies, but not ab3A-3 flies. ab3A-2 flies show significantly higher responses to the ab3A-2 ligand isopentyl acetate than ab3A-1 flies at all tested doses, and than ab3A-3 flies at the 10^-3^ dose. ab3A-3 flies show higher responses to its ligand, ethyl butanoate, than both ab3A-1 and ab3A-2 flies at 10^-4^, and ab3A-2 flies at 10^-3^. We note that all three phenotypes have similar EAG responses at higher concentrations of each odorant, which is likely because individual ORN firing rates are known to reach a maximum and plateau (1, 18). These data thus show that the changes in ab3A response between the three different phenotypes result in changes in broader antennal neuronal responses.

**Figure 1:**
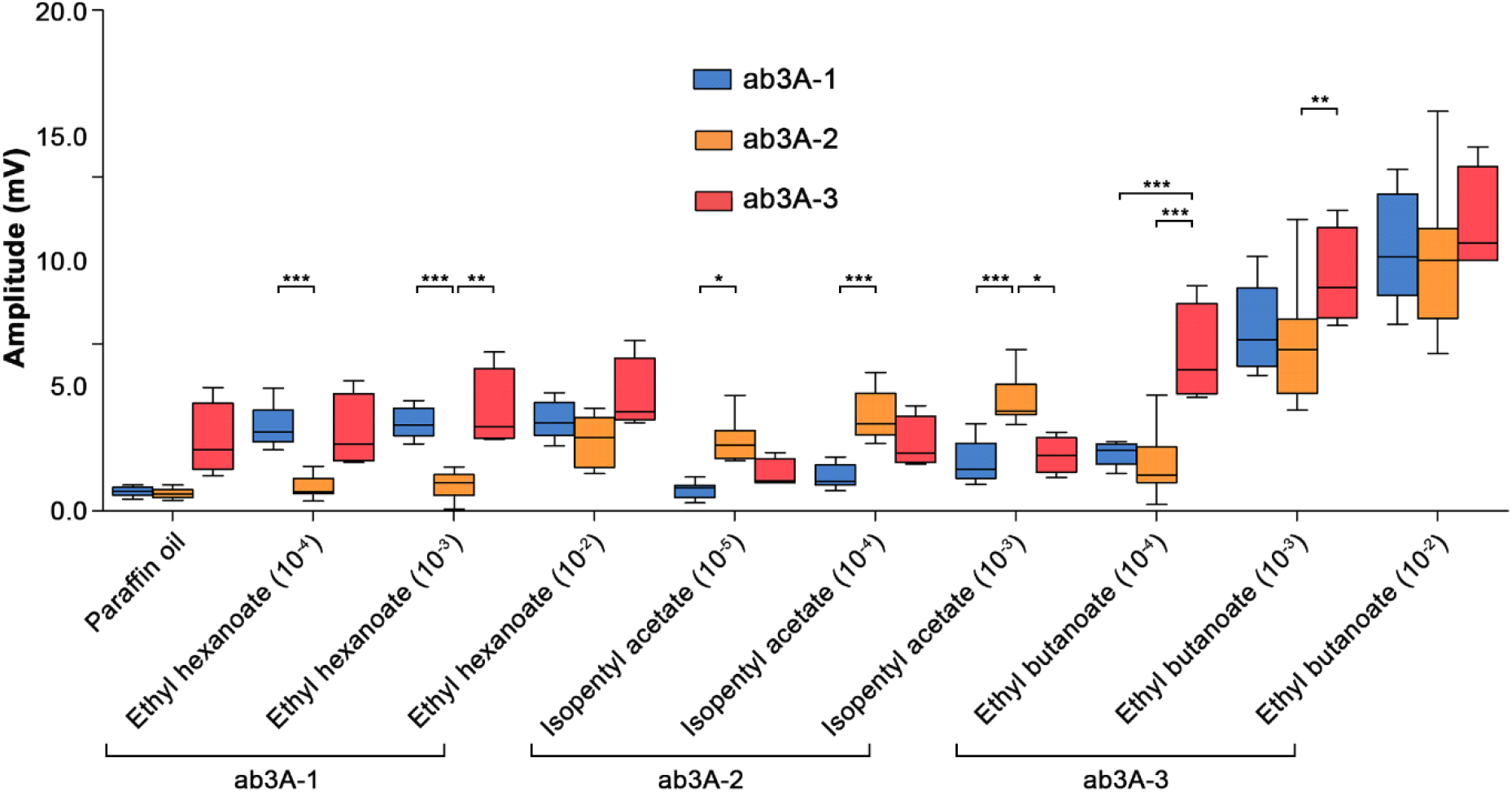
Flies with different ab3A phenotypes show variation in EAG responses. Isogenic lines with the three different ab3A phenotypes show differences in EAG amplitudes for lower concentrations of relevant odorants (2-way ANOVA with Bonferroni post-tests, *p<0.05, **p<0.01, ***p<0.001). Paraffin oil is the solvent control, doses were selected as lowest dose giving maximum response (10) ±1 dose. Data presented as median + interquartile ranges; n=9 for ab3A-1 and ab3A-2, n=4 for ab3A-3.

### Flies with the ab3A-1 and ab3A-2 phenotypes have differing behavioural preferences for ethyl hexanoate

As the changes in ab3A ORN responses alter overall detection of odorants by the antenna we proceeded to investigate whether they cause differences in olfactory behaviour responses. We were particularly interested in determining if we could identify behavioural differences between flies with the long and short *Or22* allele. As mentioned, these show latitudinal clinal distribution in Australia, with high frequencies of the short allele in the north and the long allele fully penetrant in the south. While both the ab3A-1 and ab3A-3 phenotypes are associated with the long allele, in our earlier study of this locus all isogenic lines that we derived from southern populations had the ab3A-1 phenotype. We therefore tested for behavioural differences between the ab3A-1 and ab3A-2 phenotypes. We used ethyl hexanoate as our test odorant because this odorant is not detected by other neurons at lower concentrations (8, 12, 17), and ab3A-1 and ab3A-2 flies detect this odorant differently at the level of both the individual neuron (10) and the antenna (Figure 1).

To measure olfactory preference behaviour we used a two-choice cage assay adapted from Faucher et al. (19). Flies were given a choice between two bottles with an attractive odorant added, one of which is dosed with the odorant of interest. A preference index (PI) was calculated from the number of flies in each bottle at the end of the assay. A negative PI indicates flies preferred the control bottle (C), while a positive PI indicates flies preferred the test bottle (T) with the odorant of interest added. We first tested the ethyl hexanoate preferences of flies from the ab3A-1 and ab3A-2 isogenic lines. We assessed the behavioural choices of males and females separately in order to determine if there was an effect of sex on behaviour. For both females (Figure 2A) and males (Figure 2B) we found that ab3A-2 flies were significantly more attracted to ethyl hexanoate than ab3A-1 flies (Mann Whitney U of ab3A-1 v ab3A-2: females p=0.038; males p=0.004). For both isogenic lines we observed no significant difference in preference between males and females (ab3A-1 males v females: p=0.206; ab3A-2 males v females: p=0.278). Given this, we used females only for all further experiments.

**Figure 2:**
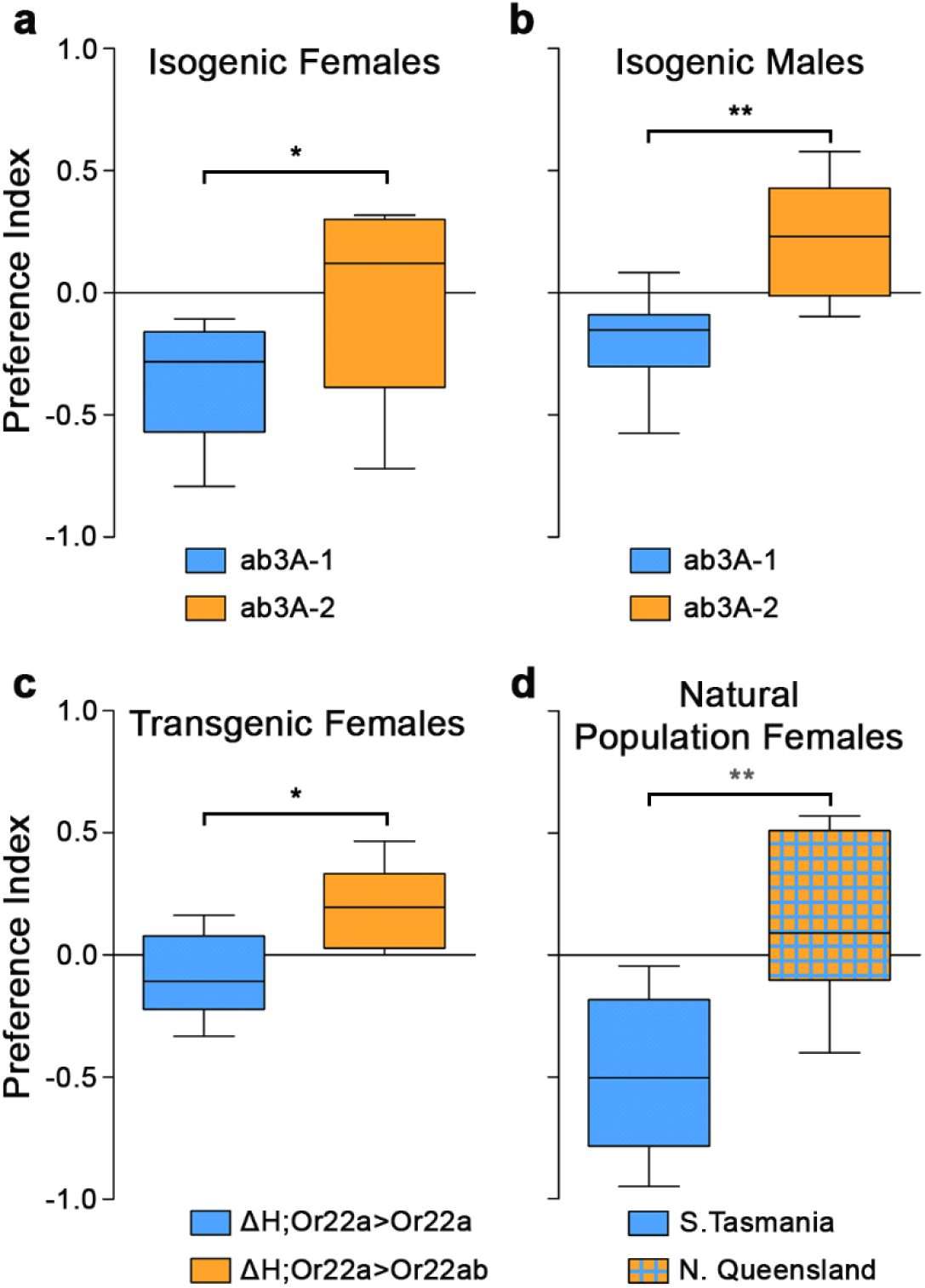
*Or22* genotype correlates with olfactory behavioural response in a two-choice cage assay. (a,b) Female and male flies from the ab3A-2 isogenic line (orange) show greater attraction to ethyl hexanoate than the same sex flies from the ab3A-1 isogenic line (blue; Mann Whitney U, a: p=0.038, b: p=0.002). Preference indices of the individual isogenic lines do not differ significantly between sexes (ab3A-1: p=0.203, ab3A-2: p=0.275). (c) Female flies expressing *Or22ab* in the empty neuron (orange) are more attracted to ethyl hexanoate than flies expressing *Or22a* in the empty neuron (blue; p=0.010). (d) Flies from a population with only the *Or22a+Or22b* allele (blue) show significantly less attraction to ethyl hexanoate than flies from a population with ~75% frequency of the *Or22ab* allele (orange with blue hatching; p=0.003). Data are presented as the median + interquartile ranges, n=8.

We note that the isogenic lines contain other differences in their genetic background aside from the genotype at the *Or22* locus, and thus these could potentially contribute to the behavioural differences observed. Therefore, to further determine if variation at the *Or22* locus leads to behavioural differences, we tested the same behavioural preference for ethyl hexanoate in females expressing either *Or22a* (ab3A-1) or *Or22ab* (ab3A-2) transgenes in the ‘empty neuron’ system (9). We found a significant difference between the preferences of flies expressing *Or22ab* and those expressing *Or22a* (p=0.014), with flies expressing *Or22ab* (ab3A-2) more attracted to ethyl hexanoate than those expressing *Or22a* (ab3A-1; Fig 2C), as would be expected if the difference observed in the isogenic lines is due to the *Or22* locus. We note that in this experiment the genetic background of the flies is not completely identical, as the background of the chromosome carrying the UAS transgene differs. Thus, while the combination of these two experiments is persuasive, we cannot definitively conclude from either alone that the behavioural differences we observe are due to the *Or22* genotype.

We therefore performed a third experiment to further support that *Or22* genotype influences attraction to ethyl hexanoate. In this case we tested attraction to ethyl hexanoate in females from two natural fly populations collected from the north and south of Australia. The population from the north (northern Queensland) was determined to have a 74.9% (±1.6%) frequency of the short allele, while the population from the south (southern Tasmania) only has the long allele present. We found that these two populations have significantly different preferences (Mann Whitney U, p=0.003), with flies from the northern population, with the high frequency of *Or22ab*, showing significantly higher attraction to ethyl hexanoate than flies from the southern population, which is fixed for the long allele (Figure 2D). The combined data from these three different experiments thus strongly suggest that flies with the *Or22ab* variant are more strongly attracted to ethyl hexanoate than are flies in which *Or22a* determines ab3A response, and thus that genotype at the *Or22* locus is associated with variation in olfactory preference behaviour.

### Flies with the ab3A-1 and ab3A-2 phenotypes have different fruit oviposition preferences

Ethyl hexanoate and other odorants detected by the ab3A neuron are known to be released from a variety of fruits, including fruits on which *D. melanogaster* flies oviposit. Along the Australian east coast there are different geographic distributions of fruits grown due to different climates. For example, tropical fruits such as banana are grown in the north, whereas fruits that grow in temperate climates, such as apple, are grown predominantly in the south. We therefore wondered if there are differences between the ab3A phenotypes in preference of fruits for oviposition, which might lead to different phenotypes being favoured in different geographic regions, and might explain the observed cline in allele frequencies along the east coast.

To test for oviposition fruit preference we designed an assay (adapted from that of Stensmyr et al. [20]) that gives females a choice between eight different fruit substrates on which to oviposit, as well as a control plate with no fruit added. Each fruit was homogenised and mixed with 1% agarose dyed blue to reduce any visual or textural influences on oviposition site choice. Based on the proportion of eggs laid on each fruit an oviposition preference index was calculated that indicates the preference of the flies to oviposit on that particular fruit. For our eight fruits we selected half that are “northern” (tropical-growing) fruits (banana, mango, pineapple and pomelo) and half that are “southern” (temperate-growing) fruits (apple, apricot, pear and strawberry). This provided the flies with a wide variety of choice, and also allowed us to determine if flies preferred to oviposit on the northern or southern fruits. We used flies from the ab3A-1 and ab3A-2 isogenic lines in this experiment, as flies expressing *Or22a* and *Or22ab* in the empty neuron have very poor fertility (likely due to other genes within the chromosomal deletion in the empty neuron background strain [21]).

We found that the two isogenic lines showed significantly different preferences for northern and southern fruits, with ab3A-2 flies laying significantly more eggs on northern than southern fruits (Figure 3A; Mann Whitney U, p=0.014), whereas ab3A-1 flies showed no preference for one group over the other. When these data were broken down to analyse oviposition preference for the individual fruits we found that, while flies from the two isogenic lines showed similar preferences for ovipositing on six of the fruits, ab3A-1 flies had a stronger preference for apricot than did ab3A-2 flies (two-way ANOVA with Bonferroni post-tests, p<0.05), and ab3A-2 flies had a stronger preference for banana than did ab3A-1 flies (p<0.001). The difference in preference for banana was highly significant, and thus to add further evidence to this finding we performed a binary oviposition preference assay where flies had to select between banana and just one other fruit. For this we used apple as a fruit on which both lines laid well (compared to control plates, one-way ANOVA with Dunnett’s multiple comparison, ab3A-1 p<0.01, ab3A-2 p<0.05), but for which we had observed no preference difference between ab3A-1 and ab3A-2 flies (two-way ANOVA with Bonferroni post-tests, apple ab3A-1 v ab3A-2, p>0.05). Supporting the initial finding, we found that ab3A-2 flies showed a strong preference for ovipositing on banana (on average laying 77% of their eggs on banana), whereas ab3A-1 flies did not discriminate between apple and banana (Figure 3B; ab3A-1 p>0.05, ab3A-2 p<0.001). These data therefore suggest that changes in ab3A phenotype correlate with altered oviposition site preference.

**Figure 3:**
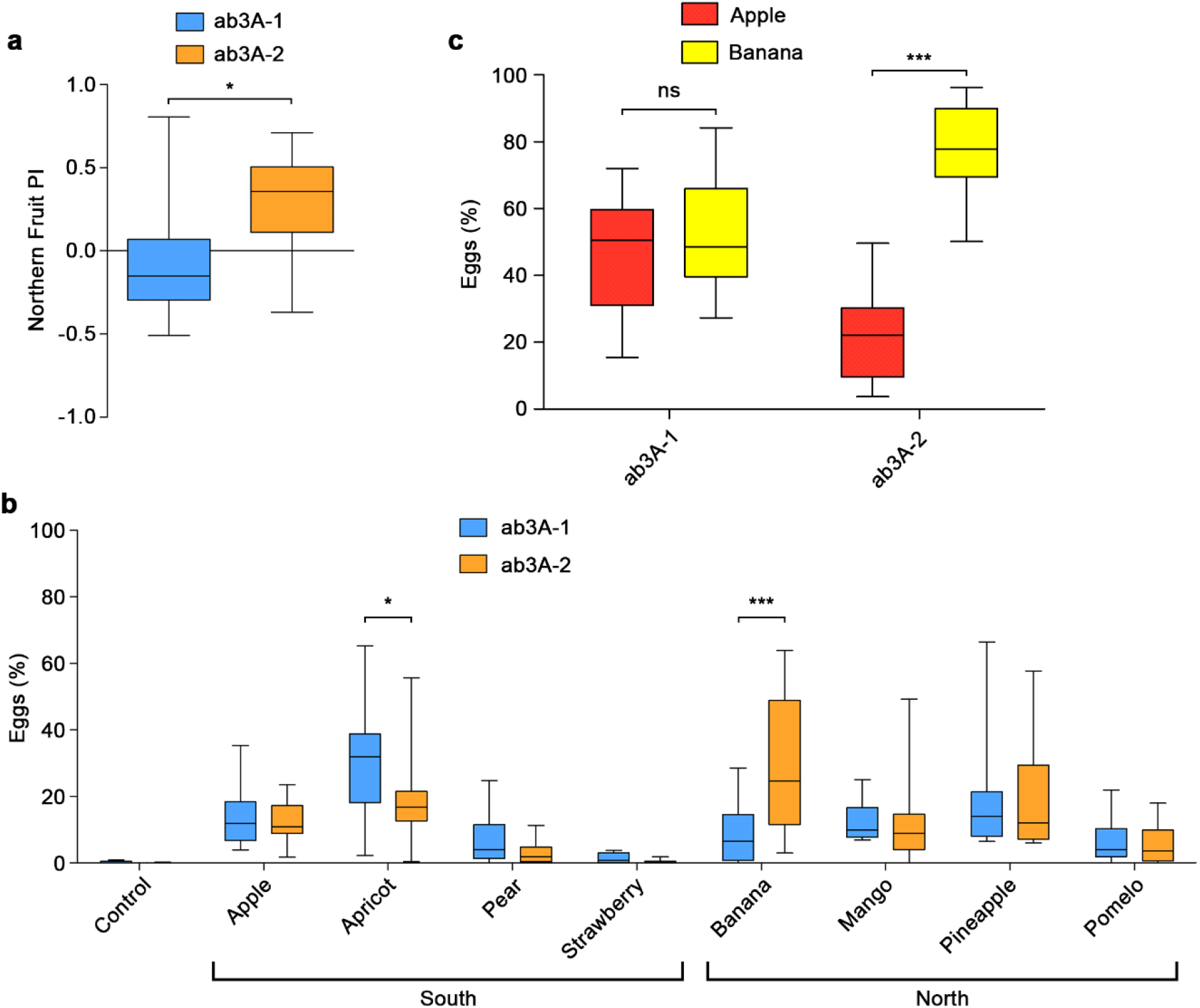
Flies with different ab3A phenotypes show differences in fruit oviposition preference. (a) When presented with four fruits typically grown in the south and four typically grown in the north, isogenic lines with the ab3A-1 (blue) and ab3A-2 (orange) phenotype showed significantly different preferences for ovipositing on northern fruits (Mann Whitney U, p=0.014). Positive index indicates a preference for northern fruits. (b) Breaking down the data from (a) for the individual fruits, choices of ab3A-1 and ab3A-2 flies differed significantly for apricot and banana (2-way ANOVA with Bonferroni post-tests; apricot p<0.05, banana p<0.001) (c) When given a choice between only apple (red) and banana (yellow) as fruit substrates, ab3A-1 flies showed no differences in oviposition site choice, whereas ab3A-2 flies strongly preferred banana (2-way ANOVA with Bonferroni post-tests; ab3A-1 p>0.05, ab3A-2 p<0.001). All data are presented as the median + interquartile range, n=12.

## Discussion

Taken together, our data suggests that ab3A response variation due to changes at the *D. melanogaster Or22* locus is associated with differences in two different olfactory-driven behaviours, and that this may then explain the cline observed in the frequencies of the long and short allele along the Australian east coast. Our finding that flies carrying the *Or22ab* allele strongly prefer to oviposit on banana is especially intriguing. Oviposition decisions are crucial to fitness because Drosophila flies lay eggs on fermenting fruits where larvae then feed on yeast. It is thus possible that the high frequency of the *Or22ab* allele found in northern Australia reflects a selective advantage in these regions where bananas are grown.

Our work contributes to a growing body of evidence that, among the many classes of ORNs, the ab3A neurons are of particular importance to the ecological specialisation of Drosophila flies. A recent study showed that a population of forest-dwelling wild *D. melanogaster* in Zimbabwe are seasonal specialists of the highly geographically restricted Marula fruit (22). The authors showed that this fruit releases high levels of the ester ethyl isovalerate, and that this is detected by ab3A neurons. They further showed that removal of ab3A neuron function reduced the ability of flies to localise marula fruit, suggesting a key role for this neuron class in this behaviour. There is also significant evidence that the ab3A neuron plays a role in host-specialisation in several other Drosophila species. *D. sechellia* are specialists of the morinda fruit, and their detection of this fruit is mediated by it releasing methyl hexanoate, which is predominantly detected by ab3A neurons (23). *D. erecta* are seasonal specialists of the *Pandanus* fruit, and detection of this fruit is predominantly due to the release of 3-methyl-2-butenyl acetate, which is also primarily detected by ab3A neurons (24). *D. suzukii* is a pest species that, unlike most other Drosophila species, prefers fresh rather than rotting fruit. The ab3A neurons of this species have been found to detect β-cyclocitral, a compound to which they are attracted that is released from leaves (25). This ligand is not detected by the ab3A neurons of *D. melanogaster*, and the authors thus proposed that this change in ligand-specificity of the ab3A neuron is one of the factors involved in the switch of *D. suzukii* from rotting to fresh fruit.

Compared to other ORN classes, the ab3A neuron response profile is unusually variable across different Drosophila species (11, 12), albeit this analysis of ORN types was not extensive. Paralleling this, in addition to showing genetic variation within *D. melanogaster*, the *Or22* locus is also unique among the *Ors* in exhibiting high levels of genetic variation and extensive levels of copy number variation across species (14). This suggests that the high level of genetic variability might provide opportunities for functional changes that contribute to the evolution of differences in olfactory-driven behaviours. While naturally-occurring changes in the sequence of some other *Or* genes have been implicated in changes in olfactory behaviour (26, 27), our data provides one of the first insights into the causative links between Or genetic variation, neuronal response variation and behavioural changes. It will be of great interest in future to determine which *Or22* genetic variants underpin shifts in host specialisation in other Drosophila species, and have thus contributed to the ability of these species to evolve and utilise different ecological niches.

## Materials and Methods

### Drosophila stocks

Four mass-bred populations of *D. melanogaster*, originally collected from Bowen, Queensland (19°58’S), Innisfail, Queensland (17°31’S), Northern Tasmania (41°S) and Southern Tasmania (43°S), were a kind gift from Carla Sgró, Monash University (28). The allelic frequencies of the long and short alleles in the northern Queensland population were found by taking 14 samples of 56 to 92 flies and individually PCR genotyping all flies in the sample using GoTaq (Promega). Primer sequences: Or22ab-F 5’ GCA AGT TTT TTC CCC ACA TT 3’; Or22ab-R 5’ ACC CCA TGA GAA TGA CTT CG 3’; amplicon is 336bp; Or22a&b-F 5’ GCA GTT TTT CGC AAA GGA AG 3’; Or22a&b-R 5’ AAA GTT TTC CGG GAA TGT CA 3’; amplicon is 639bp. Isogenic lines were derived as described in Shaw et al. (10). All isogenic lines were maintained on standard wheat-based media at 22°C. To drive expression of olfactory receptor transgenes in the ab3A neuron we used flies carrying Δ*Halo*, a small deletion on the second chromosome that removes the *Or22* locus, combined with an *Or22a*-promoter construct (13). The *w;ΔHalo/CyO;P[Or22a-Gal4]* and *w;ΔHalo/CyO;P[UAS-Or22a]*, stocks were obtained from John Carlson, Yale University. *P[UAS-Or22ab]* stock is from Shaw et al. (10). All crosses were performed at 25°C.

### EAG recordings

Electroantennograms (EAGs) were recorded as per Tom et al. (29). Briefly, a single fly was immobilised and a reference electrode inserted into the eye and a recording electrode placed on the surface of the antenna. Changes in voltage (mV) in response to 1s stimulations with odorants were amplified using an active probe and a serial-IDAC amplifier (Syntech). EAGs represent the summed activity of a population of ORNs. Odour stimulation was by injecting volatiles from 5mL syringes into an airstream blown over the preparation. All odorants were at highest available purity (>98%, Sigma-Aldrich) and dissolved in paraffin oil to reach the specified dilution.

### Two-Choice cage assay

The two choice cage assay was adapted from Faucher et al. (19). A 305×305×305mm cage was fitted with two 66mm funnels placed 130mm apart leading into 250mL glass bottle traps. Bottles were filled with 40mL solutions of 50% (vol/vol) apple cider vinegar (Cornwall’s) to attract flies. The test odorant was added to one of the traps (10-4 vol/vol for ethyl hexanoate). When testing isogenic lines one hundred flies were starved on 1% agarose for 24 hours before being released into the cage. When testing the flies expressing either *Or22a* or *Or22ab* in empty ab3A neurons between 50 and 80 flies were tested, due to the poor health of these flies. The cage was surrounded by an open-top black box to minimise directional visual stimuli and placed in a fume hood to provide ventilation. The assay was run for 17 hours under natural light conditions, such that the dawn and dusk activity periods were captured. The side that contained the test odorant was alternated between every repetition. A preference index was calculated as (T-C)/(T+C) where T is the number of flies in the test odorant-containing bottle and C is the total number of flies in the control bottle. The preference indices were then compared for significance using a Mann Whitney U test.

### Oviposition behaviour assay

The oviposition assay was adapted from Stensmyr et al. (20). We tested eight fruits representing those commonly grown in tropical northern Australia (banana, mango, pineapple and pomelo) and in temperate southern Australia (apple, apricot, pear and strawberry). Fruits were purchased from a fruit and vegetable market, aged for three days at room temperature and then homogenised and frozen. One gram of thawed homogenised fruit was added to the lid of a 35mm tissue culture plate (Sarstedt) with two drops of blue food dye and 4mL of 1% agarose minimising both visual and textural differences. Once cooled and set, the plates were placed into a 235×235×120mm plastic box with a damp sponge for humidity, a lid with holes for ventilation and left for 24 hours at room temperature (22°C) before flies were added. Nine plates were placed in a 3×3 grid pattern; one for each of the eight fruits in a pseudorandomized distribution and a control plate containing water in the middle position. When only apple and banana were tested the two fruits were distributed in a regular pattern over the eight plates, with the fruit in the corner plate alternating and the control plate remaining the same. Virgin females and males were kept separately for three days and then crossed. After two days 40 fertilized females were placed in a small cage with a food plate coated with yeast paste to induce laying. After two days laying on food plates, 30 females were gently sedated by cooling and released in the box. The assay was run for 24 hours and the number of eggs laid on each plate was counted to calculate the proportion of eggs laid on each fruit out of the total.

## Acknowledgements

We thank the Australian *Drosophila* Biomedical Research Facility (OzDros) and Jyotika Taneja de Bruyne for technical support, the Bloomington *Drosophila* Stock Centre (BDSC), John Carlson and Carla Sgró for providing fly stocks, and Ary Hoffman, Stephen McKechnie, Damian Dowling, Charles Robin and Tim Connallon for valuable discussions.

## Author Contributions

C.G.W. and M.deB. conceived the experiments, interpreted the data and co-led the work. K.H.S. conceived the experiments, interpreted the data and performed the experiments. C.I.D. performed experiments. A.A. and T.K.J. interpreted the data. K.H.S. and C.G.W. wrote the paper.

---

**Supplementary Table 1:**
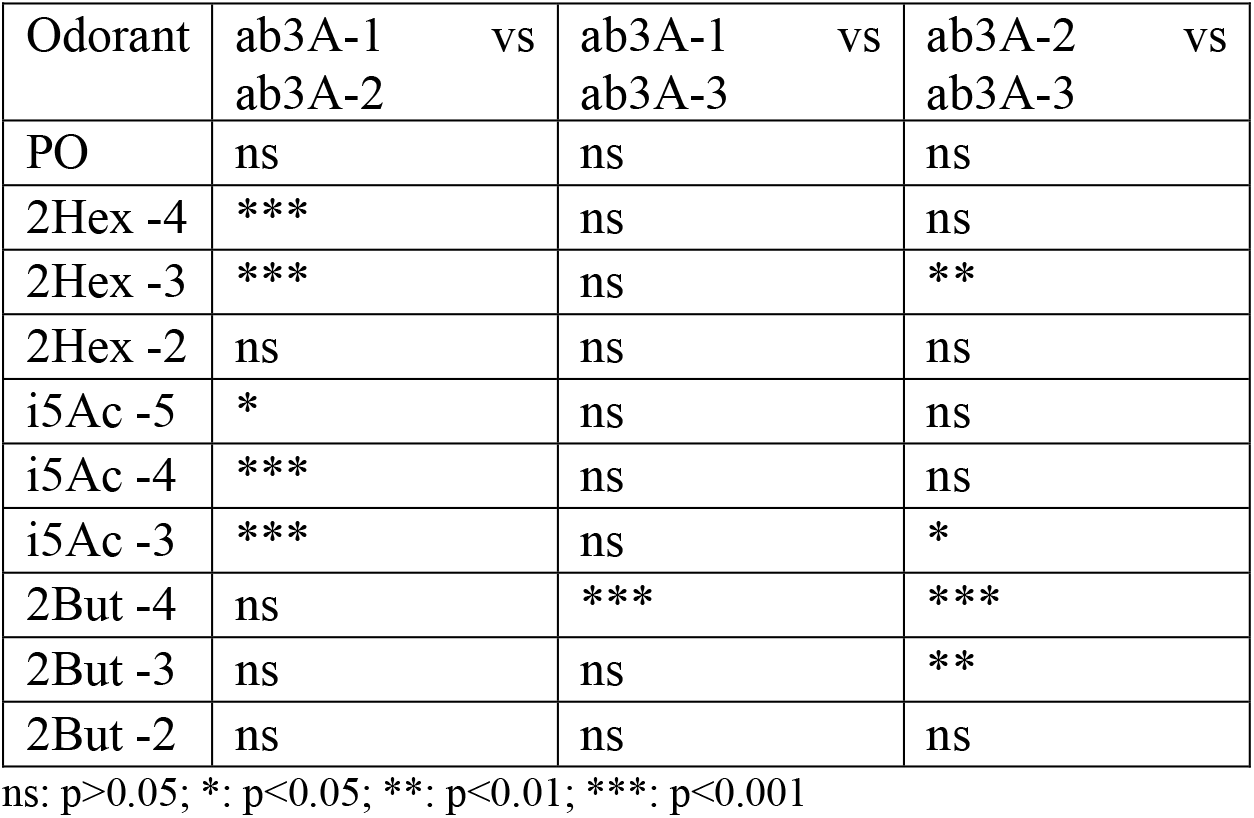
2-way ANOVA with Bonferroni post-tests of data from Figure 1

